# Longitudinal Proteomic Profiling Defines Robust Molecular Subtypes Underlying the Heterogeneity of Parkinson’s Disease

**DOI:** 10.1101/2025.11.27.691036

**Authors:** Rashmi P Maurya, Buddhiprabha Erabadda, Sacha E Gandhi, Hanjun Zhao, Natacha Loison, Raquel Real, Huw Morris, Alejo Nevado-Holgado, Donald G Grosset, Laura M Winchester

## Abstract

Heterogeneity of Parkinson’s disease (PD) pathology is a barrier to developing therapeutics and understanding progression and prognosis. High throughput proteomic measures can be used to better interpret PD pathophysiology and generate clusters to define disease subtypes. Identification of subtypes related to clinical phenotypes will help researchers understand PD progression.

We analyzed longitudinal proteomic data from the Tracking Parkinson’s Cohort, consisting of recent-onset PD patients across 72 UK sites. 794 patients were measured on the Somalogic platform (7596 proteins) for three time points. Weighted Gene Co-Expression Network Analysis (WGCNA) at each time point revealed consistent protein co-expression modules. Two modules were strongly preserved across all three time points and in validation in the Global Neurodegenerative Proteomics Consortia (GNPC) datasets. The brown module was enriched for metabolite pathways and the blue module with cellular signaling pathways and associated with quality of life scores. Conversely, the smaller red module had distinct cognitive function phenotypes and changed protein expression between visit time points.

Using detailed characterization of proteomic clusters we have provided a comprehensive view of PD progression offering deeper insights into the conservation of proteomic expression, suggesting new module subsets and providing candidate target proteins for further study.

## Introduction

The heterogeneity of Parkinson’s disease (PD) is one of the main barriers to successful identification of therapeutic drugs and robust biomarkers^1^. Progression of PD varies dramatically between individuals. Onset and progression of disease traits show heterogeneity in rates of change and severity^2^. Resolving subtypes of disease pathology will enable researchers to better understand differences in mechanism and develop precision treatments. The current SynNeurGe classification frameworks suggest Alpha-synuclein, neurodegeneration and pathogenic genes^3^ define subtypes; however, these standards are not universally accepted and further work is needed to establish the underlying physiopathology and mechanism of these groupings^4^. Multi-omics assays offer measures from the pathways and processes responsible for disease pathogenicity and, as such, are potential candidate markers for detecting these groups and subtypes. Here, we use protein expression to define common patient and pathology groupings. The primary goal of this study (Figure 1) was to define proteomic subtypes to understand the heterogeneity of Parkinson’s pathophysiology with the aim of understanding whether there exists an underlying biological signature of mechanism in distinct disease phenotypes. By running unsupervised clustering approaches we have detected common modules of expression. Uniquely we have used longitudinal data to compare these groupings during disease progression. Despite the complexities of disease progression we describe robust proteomic modules with insights about mechanism and subtypes of pathology.

**Figure 1.**
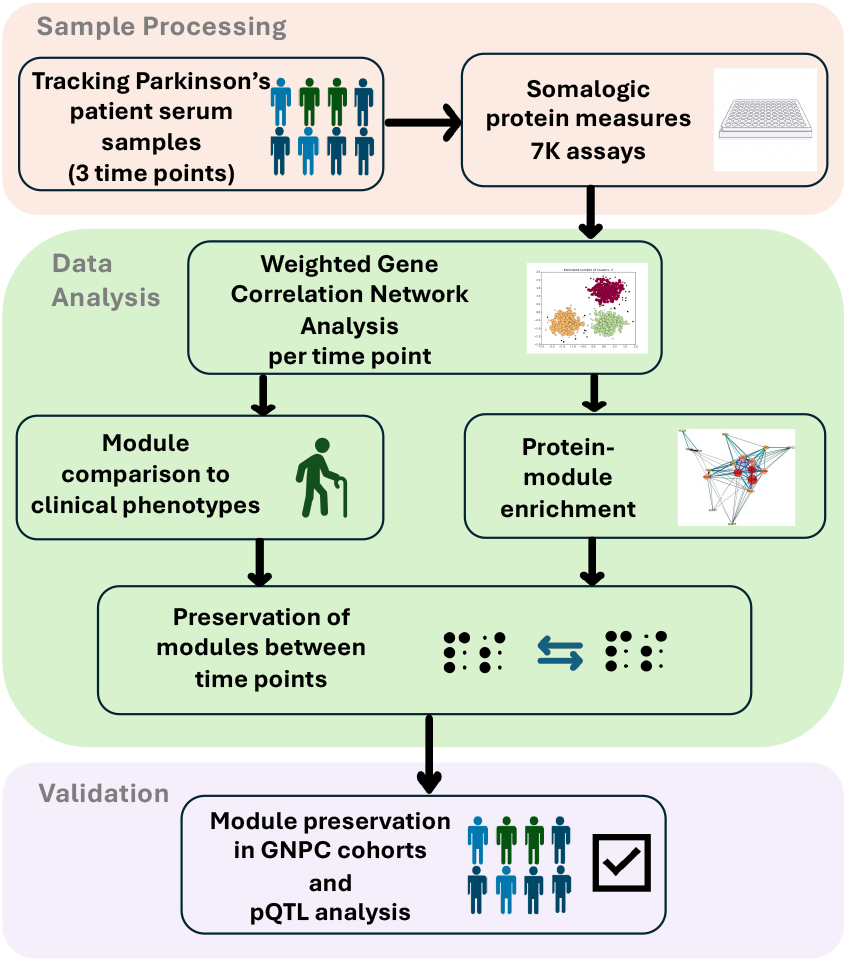
Overview of the study design and analytic approaches.

## Results

### Longitudinal demographics measures for the Tracking Parkinson’s cohort

A total of 757 participants from the Tracking Parkinson’s (TP) cohort were investigated across three time-points. The participant characteristics for these visits are described in Table 1. There were more males (66.1%) than females (33.7%) in the cohort, with a mean age of 66.3 years at baseline. Disease severity progressed substantially over time, as evidenced by significant increases in Movement Disorders Society Unified Parkinson’s Disease Rating Scale (MDS-UPDRS) III total scores (Visit 1: 21.06±11.24 vs Visit 3: 30.27±15.06; F(2,1300.8)=182.63, p<0.001, Cohen’s d=1.06) and Hoehn and Yahr staging. Medication received increased substantially over the study period, as reflected by significant increases in Levodopa equivalent daily dose (LEDD) across visits (Visit 1: 275.97±195.44 mg vs Visit 3: 577.96±309.32 mg; F(2,1529.0)=718.83, p<0.001). Notably, MoCA total scores showed a modest but significant decline over time (Visit 1: 25.59±3.29 vs Visit 3: 25.20±4.04; F(2,1368.8)=14.58, p<0.001), with post-hoc pairwise comparisons revealing a significant decrease between Visit 2 and Visit 3 (mean difference: 0.59 points, p<0.001, Cohen’s d=0.29). Individually, each time point mean score was indicative of mild cognitive impairment.

**Table 1.** Comparison of Tracking Parkinson’s cohort demographics between visits.

| Characteristic | Visit 1 | Visit 2 | Visit 3 |
| --- | --- | --- | --- |
| Total samples | 757 | 757 | 757 |
| Mean age at visit [SD] (IQR) | 66.3 [8.6] (61.3–72.1) | 67.9 [8.5] (62.8–73.7) | 69.5 [8.5] (64.4–75.2) |
| *Sex (%)* |  |  |  |
| Female | 255 (33.7) | 255 (33.7) | 255 (33.7) |
| Male | 500 (66.1) | 500 (66.1) | 500 (66.1) |
| Mean disease duration [SD] (IQR) | 1.3 [0.9] (0.5–2.0) | 2.8 [0.9] (2.0–3.6) | 4.4 [1.0] (3.6–5.1) |
| Mean LEDD total [SD] (IQR) | 276.5 [196.5] (100.0–400.0) | 431.6 [266.2] (300.0–550.0) | 580.5 [311.0] (400.0–720.0) |
| Mean UPDRS III [SD] (IQR) | 21.1 [11.2] (13.0–27.0) | 26.4 [13.1] (17.0–34.0) | 30.2 [15.1] (19.0–40.0) |
| Mean MoCA total [SD] (IQR) | 25.6 [3.3] (24.0–28.0) | 25.8 [3.5] (24.0–28.0) | 25.2 [4.0] (23.0–28.0) |
| *Hoehn & Yahr stage (%)* |  |  |  |
| Stage 1 | 270 (35.9) | 135 (18.0) | 94 (12.6) |
| Stage 1.5 | 135 (18.0) | 122 (16.2) | 87 (11.7) |
| Stage 2 | 239 (31.8) | 305 (40.6) | 300 (40.2) |
| Stage 2.5 | 77 (10.2) | 124 (16.5) | 134 (18.0) |
| Stage 3 | 30 (4.0) | 59 (7.9) | 106 (14.2) |
| Stage 4 | 1 (0.1) | 6 (0.8) | 19 (2.5) |
| Stage 5 | 0 | 0 | 6 (0.8) |

LEDD = Levodopa equivalent daily dosage; UPDRS = Unified Parkinson's Disease Rating Scale; MoCA = Montreal Cognitive Assessment.

### Cross-sectional Clustering Analysis Reveals Distinct Functional Modules

Using clustering, we sought to identify protein co-expression modules in a cross-sectional analysis. We used 7288 proteins from the TP cohort as features in the clustering algorithm as explained in the methods section and independently analyzed visits 1, 2, and 3 (denoted as V1, V2, and V3, respectively, in the subsequent text) to understand how the clusters are preserved across the visits. Finally, we carried out pathway, tissue, and cell-type enrichment analyses for each cluster to identify the biological role of proteins in the clusters.

We performed cross-sectional clustering using Weighted Gene Co-expression Network Analysis (WGCNA)^5^ to identify distinct protein co-expression modules for the first three visits of the TP Cohort. Excluding the *grey* (unassigned) module, we observed seven common modules across all visits and one additional module in V3 (Supplementary Table 1 for detailed protein module assignments for all visits). We observed substantial overlap between consensus WGCNA modules and those from cross sectional visit-specific modules. The module sizes (Table 2) were very similar across approaches and the corresponding modules exhibited concordant pathway enrichment profiles (Supplementary Fig 1).

**Table 2.** Protein co-expression module sizes for WGCNA networks constructed separately at each visit and for the consensus WGCNA network.

| Module | Cross-sectional WGCNA networks |  |  | Consensus WGCNA network |
| --- | --- | --- | --- | --- |
|  | Visit 1 | Visit 2 | Visit 3 | Size (genes) |
| Turquoise | 5281 | 5203 | 5133 | 5369 |
| Blue | 614 | 694 | 777 | 704 |
| Brown | 248 | 229 | 202 | 217 |
| Yellow | 51 | 52 | 68 | 52 |
| Green | 39 | 28 | 45 | 41 |
| Black | 25 | 52 | 25 | 29 |
| Red | 29 | 33 | 23 | 31 |
| Magenta | — | — | 30 | — |
| Grey<sup>a</sup> | 1001 | 997 | 985 | 845 |

<sup>a</sup> "Grey" denotes unassigned genes in WGCNA. Em dashes indicate the module did not appear at that visit or in the consensus.

### Exploring protein modules using clinical phenotypes

To understand the relevance of the modules to potential clinical outcomes we compared each module to a panel of disease phenotypes representing different features of PD. Correlation analysis between module eigen-proteins and clinical phenotypes for each visit, revealed several module associations that remained consistent across visits for the “age at visit” phenotype. However, there are multiple differences between the correlations at V1 and V3 (Figure 2). For example, the blue and green modules are correlated with the quality of life scores (PDQ8 and EQ5D) at V1 only, and despite being one of the least stable, the red module shows stronger correlations to cognitive phenotypes (MoCA and semantic fluency), and REM Sleep Behavior Disorder (RBD total) at V3. We saw the strongest correlations between the largest turquoise module and the clinical outcomes at both time points.

**Figure 2.**
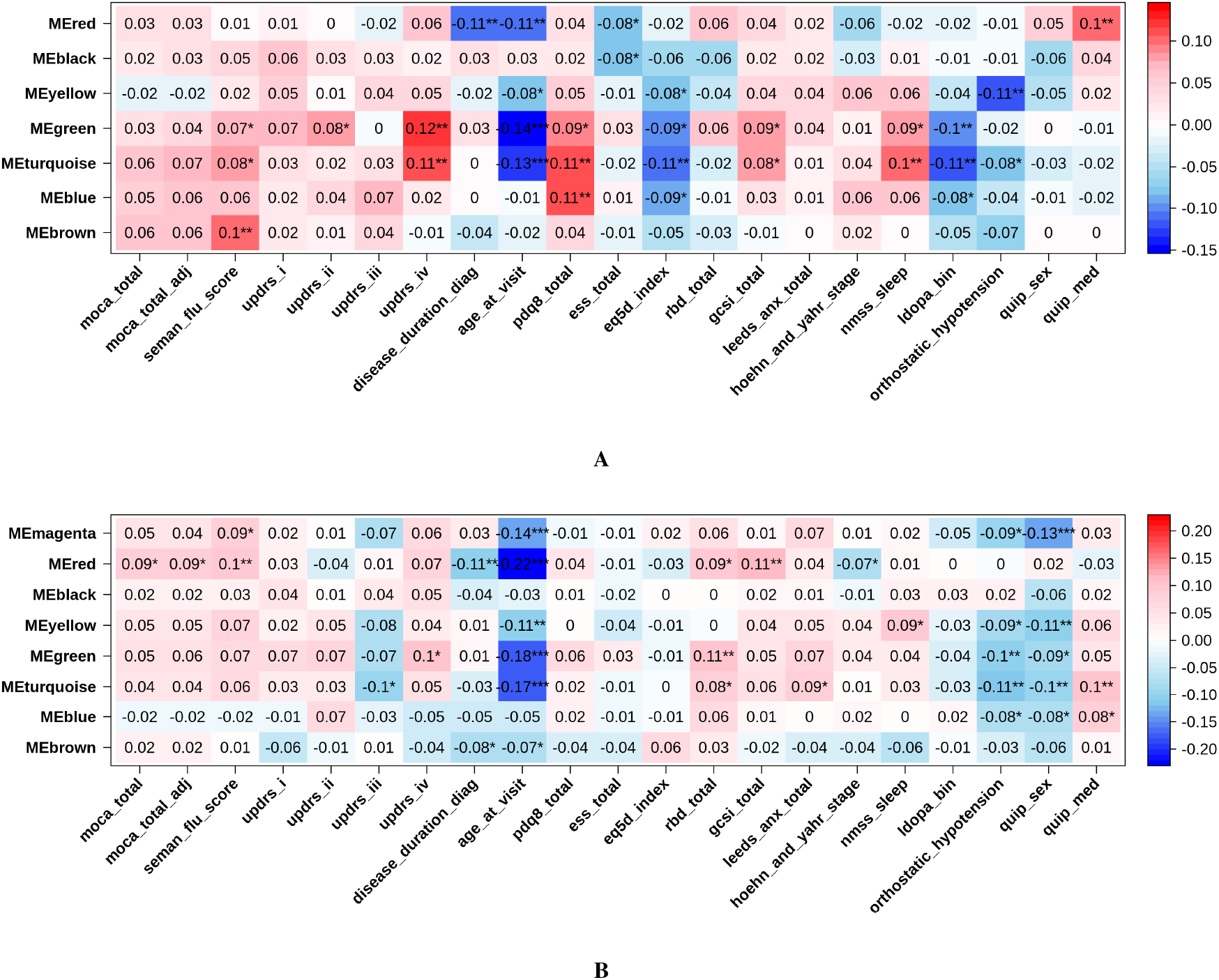
Pearson correlation associations between protein co-expression modules and clinical phenotypes at Visit 1 and Visit 3 respectively. Color intensity reflects the strength and direction of the correlation (blue = negative, red = positive). Abbreviations: seman_flu_score = semantic fluency score, a higher score corresponds with better cognition; pdq8_total = Parkinson’s Disease Quality of Life 8-item version, higher score corresponds to worse quality of life; ess_total = Epworth sleepiness scale - measure of daytime sleepiness, higher score corresponds to more daytime sleepiness; eq5d_index = EuroQol health states - quality of life index, higher score corresponds with better quality of life; rbd_total = REM Sleep behavior disorder, higher score corresponds to greater REM sleep behavior problems; gcsi_total = Gastroparesis Cardinal Symptom Scale, a higher score indicates more severe symptoms; leeds_anx_total = Leeds anxiety scale, a higher score corresponds with more anxiety; hoehn_and_yahr_stage = Hoehn and Yahr stage - Parkinson’s disease severity, Stage 0 means no signs of disease and Stage 5 means wheelchair-bound or bedridden unless aided; nmss_sleep = Non-Motor Symptoms Scale for Parkinson’s Disease (NMSS) for sleep; ldopa_bin = Patient’s Levodopa treatment status; quip_sex and quip_med = Questionnaire for Impulsive-Compulsive Disorders in Parkinson’s Disease (QUIP) for sex and medication

### Protein pathology of modules implicates known and novel pathways and mechanisms

Pathway enrichment analysis using the KEGG library for modules across visits revealed five modules that are functionally significant for the first, second, and third visits (Figure 3A, Supplementary Table 2). The turquoise module across all the visits was enriched for one pathway – cytokine-cytokine receptor interaction pathway, which is closely related to inflammation. The brown and red modules were enriched for many common metabolism related pathways. The smallest red module has stronger enrichment at the initial stages of the disease perhaps reflecting initial disease changes. The blue module is enriched for dopaminergic synapse and a number of signaling pathways including T-cell and neurotrophin. The black module includes the complement and coagulation pathway in addition to disease and infection pathways. To understand whether modules represented particular peripheral changes we ran a secondary enrichment analysis comparing organ expression libraries. Analysis shown in Figure 3B shows a difference between expression whereby blue (signaling) and brown (metabolic) modules were enriched for brain cell expression libraries and red, black and turquoise modules were enriched for whole body gene sets such as intestines and liver. Full tissue enrichment results are provided in Supplementary Table 3. A focused analysis to look at brain cell type enrichment showed an overlap between modules enriched for inflammatory pathways (turquoise, blue and black) and the cell type microglia (Figure 3C). The brown module, which has a significant enrichment for metabolic function pathways, is enriched for oligodendrocytes across the three time points.

**Figure 3.**
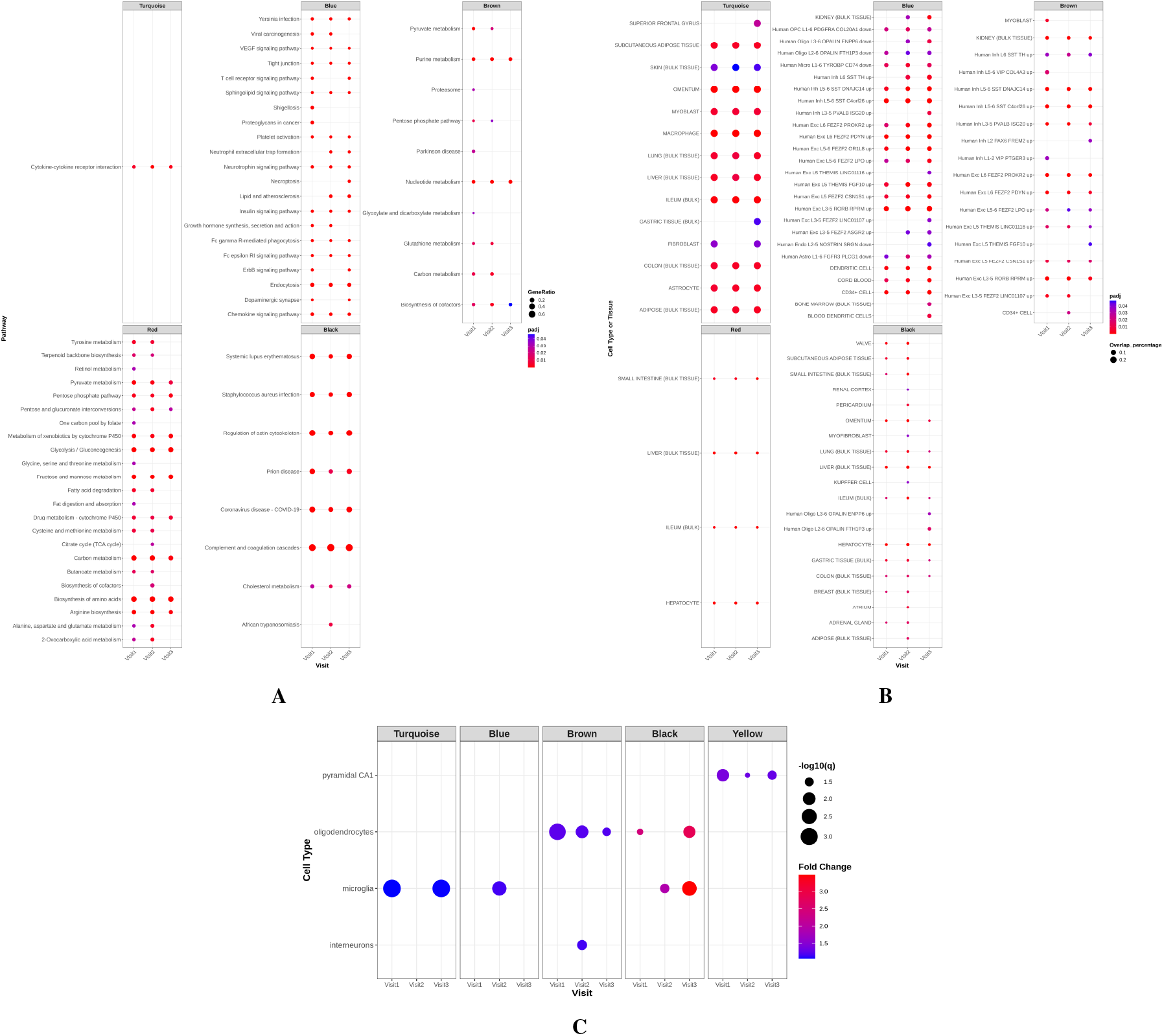
Protein module pathway enrichment in modules across visits. (A) Dot plot showing up to top 25 enriched KEGG biological pathways per module and time point. Dot size indicates gene ratio, and color intensity reflects statistical significance. (B) Dot plot showing cell type and tissue enrichment for co-expression modules across visits. Abbreviations : OPC = oligodendrocyte precursor cell; Oligo = oligodendrocyte; Exc = excitatory neuron; Inh = inhibitory interneuron; VIP = vasoactive intestinal peptide interneuron; SST = somatostatin interneuron; PVALB = parvalbumin interneuron; Micro = microglia; Astro = astrocyte; Endo = endothelial cell. Terms ending in “up” or “down” indicate the direction of differential expression in the reference dataset used by enrichR R package (C) Dot plot showing cell type enrichment for co-expression modules across visits.

Figure 4 shows the network developed between proteins and pathways of different modules reflecting the complexity of pathway membership for individual proteins. We can see the complex network for the larger blue module, which also shares proteins with the PD pathway from the brown module. There are also common links between the red and brown modules, both enriched for different metabolism pathways although we can also see the increased preservation between visits on the brown module shown by the three visit pie chart in the centre of each protein.

**Figure 4.**
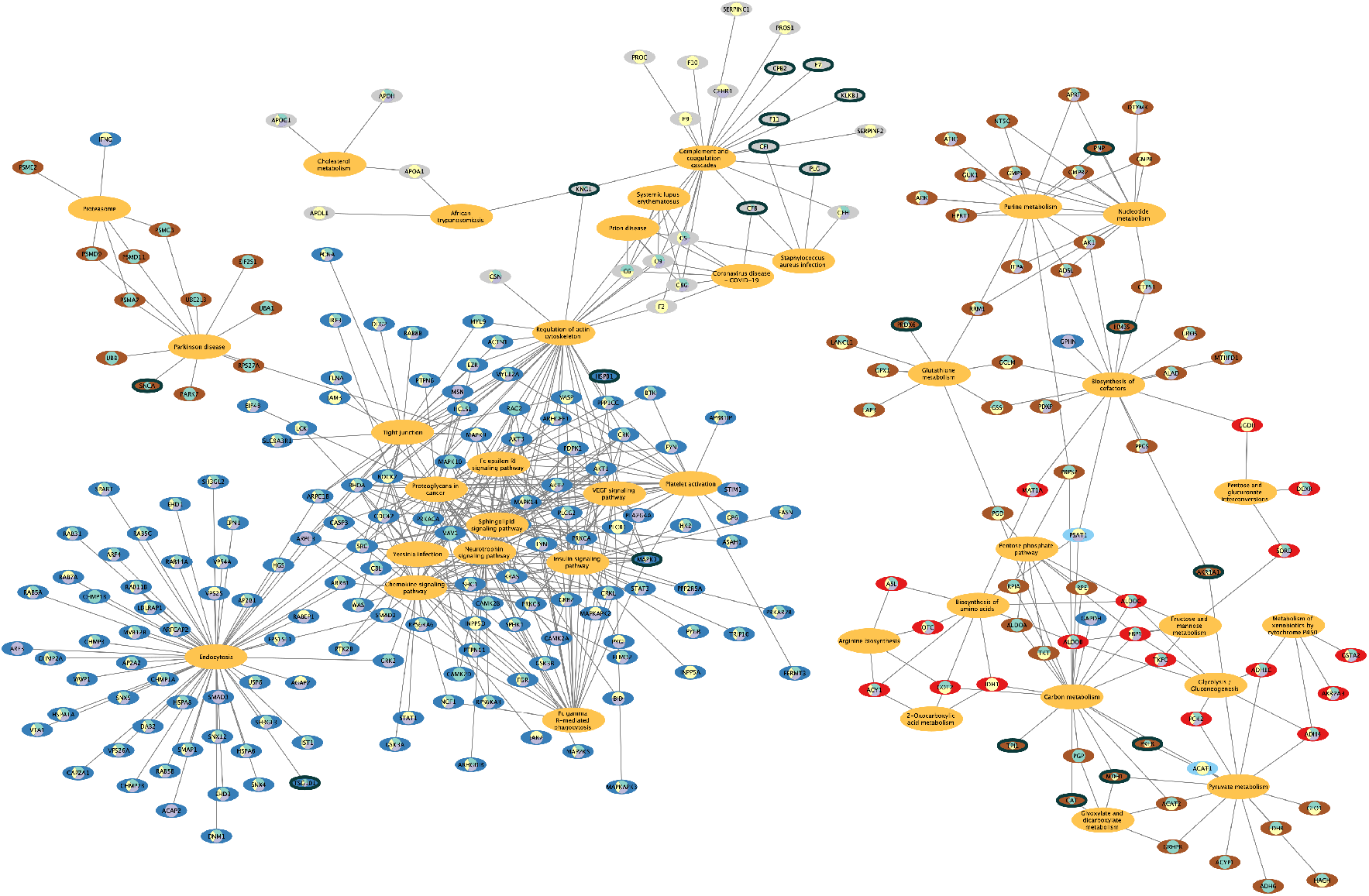
Protein-Pathway Network. The network visualizes relationships between proteins (labeled by gene names) and up to top 25 KEGG pathways (in *yellow*) based on the adjusted p-value for each preserved protein co-expression module of interest. Protein node colour indicates module membership where *grey* nodes correspond to *black* module. Node pie charts indicate the presence of each protein across three visits: visit 1 (*blue*), visit 2 (*yellow*), and visit 3 (*purple*). Nodes with black borders represent proteins with a significant (*p <* 0.05) pQTL signal.

### Module preservation analysis shows stability of modules over time

The module preservation analysis using WGCNA showed remarkable stability of modules across visits (Figure 5A). Specifically, we compared the preservation across three settings of reference (ref) and test: V1 (ref) vs V2 (test), V1 (ref) vs V3 (test), and V2 (ref) vs V3 (test). We observed that the larger modules (*turquoise, blue*, and *brown*), consistently showed evidence of strong preservation (*Z*_*summary*_ *>* 10) while smaller modules (*yellow, green, red*, and *black*) showed low to moderate evidence of preservation (2 < *Z*_*summary*_ < 10). We checked a second composite statistic (medianRank) obtained from the same analysis, which shows no dependence on module size to confirm the initial preservation scores. The medianRank statistic can lead to biologically meaningful comparison of relative preservation among modules, where lower values indicate stronger preservation^6^. This revealed that the largest turquoise and even the grey module (unclustered protein module) had higher ranks indicating lower preservation (medianRank = 6 - 9, Supplementary figure S2A). The blue (signaling) and brown (metabolite) showed high preservation in both composite statistics (medianRank = 2 - 5).

**Figure 5.**
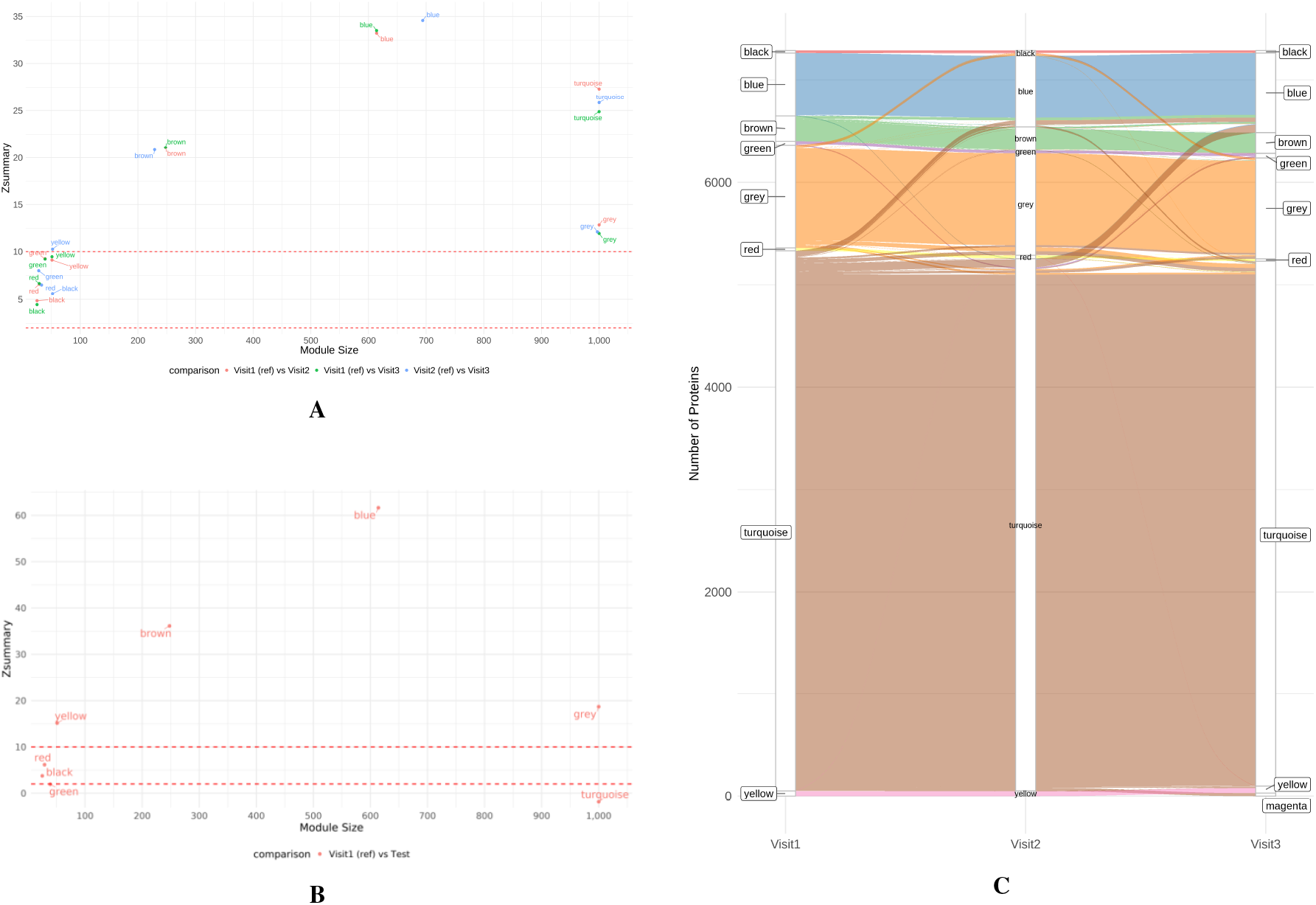
Protein module preservation and transitions across visits. (A) Preservation of co-expression protein modules across the three visits. An *Z*_*summary*_ *<* 2 indicates no evidence of module preservation, 2 *< Z*_*summary*_ *<* 10 indicates weak to moderate evidence of module preservation, and *Z*_*summary*_ *>* 10 indicates strong evidence of module preservation. (B) Preservation of protein modules in the GNPC cohort using 440 PD cases. (C) Sankey flow plot showing how proteins move between co-expression modules across visits.

The Sankey plot, which tracks protein membership within modules across visits, shows that most of the modules are temporally stable, with proteins largely remaining in the same modules across the three visits. (Figure 5C). This differential pattern of module stability suggests that specific biological processes/pathways are likely consistent and maintained throughout the course of the disease, while others undergo changes.

### Identification of cis Protein Quantitative Trait Loci (pQTLs)

The pQTL analysis identified 309 proteins (Supplementary Table 4) with at least one *cis* signal, where *cis* was defined as variants within ± 1Mb of the corresponding gene. These proteins were analyzed against the WGCNA module membership for each visit to identify proteins that are genetically influenced and the counts are provided in Supplementary Table 5. To further prioritize relevant protein signals we used module hub proteins significantly associated with *cis* pQTL hits (Supplementary Table 6). The blue (signaling) module consistently pointed to cellular stress and survival signaling. MAPK3 (also known as ERK1) was identified in all three visits. This suggests that ongoing cellular stress responses are a stable component of the molecular landscape of PD. The black (infection and disease) module consistently included proteins from the complement and contact systems, key branches of innate immunity and coagulation. Across the visits, CFI, and KNG1 were recurring members. In Visit 2, this module also contained significant pQTLs for coagulation factors like F7. This stable signature suggests that neuroinflammation and potentially vascular-related disruptions are sustained features of the disease process.

### Validation shows modules are similarly preserved in external cohorts

Using the Global Neurodegeneration Proteomic Consortium (GNPC) replication cohort, PD cases were selected to demonstrate module performance in independent data cohorts with alternative blood sample matrix (plasma). This showed that the modules are preserved with similar evidence compared to the discovery cohort (Figure 5B, Supplementary Figure 2B). Specifically, using baseline modules from TP compared to cross-sectional data from the replication cohort, two of the core modules were validated. The *blue* and *brown* modules were strongly preserved with *z*_*summary*_ *>* 10 (medianRank = 3), while *red, black*, and *green* modules showed weak to moderate evidence of module preservation. Interestingly, the turquoise module, which was highly preserved in our discovery cohort, was not observed in the replication cohort. This suggests that the biological processes represented by the turquoise module are highly dynamic. The module’s strong and stable activation appears to be a feature of our discovery cohort which is absent in the replication cohort. This could point to heterogeneity in clinical sample collection between cohorts but could also relate to the difference in blood matrix and protein aptamer performance on this module subset.

## Methods

### Patient Cohort

The Tracking Parkinson’s cohort (Parkinson’s Repository of Biosamples and Networked Datasets PRoBaND) is a multi-center, observational, prospective study of Parkinson’s disease, with blood samples collected across 72 UK sites from 2012 until 2022^7^. It follows recent onset Parkinson’s disease patients (diagnosed within the last three years). The samples were initially collected every 6 months for 3 years and then every 18 months for up to 7 years. The Somascan assay was used to measure 7596 proteins from blood plasma for 1918 individuals at up to 5 time points. We used a subset of this data for this study to ensure robust analysis. This included common patients (ie, 794 patients) across the first 3 time points (baseline visit, 18 months and 36 months visits). Motor, non-motor, and quality of life assessments were performed using validated scales as described by Malek et al^7^. These were repeated at six month intervals or at each additional visit.

### Ethics and Data Availability

The Tracking Parkinson’s study is carried out in accordance with the Declaration of Helsinki. Patient consent was obtained and ethics approval was made by the West of Scotland Ethics Committee. Proteomics data is available on the AD work-bench (https://discover.alzheimersdata.org) via application to the Global Neurodegeneration Proteomics Consortium (GNPC) https://www.neuroproteome.org/

### Proteomic Profiling

For each sample, protein levels were measured using a Slow Off-rate Modified Aptamer (SOMAmer)–based capture array (SomaLogic, Inc., Boulder, CO). Protein concentration measurements were obtained from Relative Fluorescent Units (RFU) that had been processed using SomaLogic’s proprietary normalization pipeline to reduce inter-plate variability.

### Quality Control and Data Preprocessing

From the initial data set of 2382 samples across three visits and 7596 proteins, we first filtered the proteins to include only 7288 human proteins. Our initial data exploratory analysis with principal components revealed the presence of technical batch effects arising from different sites. To resolve this, the data was first *log*2 transformed and then corrected for site-related batch effects using the sva package^8^ in R. Finally, we removed samples with missing sex and age information from the analysis resulting in 771 common patients across visits.

For the pQTL analysis, samples were genotyped using the Illumina HumanCoreExome array. We first excluded samples with >2% missingness (PLINK −− mind). We also excluded samples with undefined sex or where the reported sex did not match the genetically inferred sex (PLINK −− check-sex 0.2 0.7), duplicates and related individuals (PLINK −− genome −− max 0.125) and heterozygosity outliers (PLINK −− het; |Z| > 2 SD). The genotypes used in the analysis were imputed against the HRC panel (v.r1.1 2016) in the Michigan Imputation Server using Minimac4 (v.1.0.0). Post imputation, variants with imputation score R2 ≤ 0.3, missingness >5% (PLINK −− geno) and MAF <1% (PLINK −− maf) were excluded. We then matched the genotyped samples to clinical records to select samples with a matching record, resulting in 799 cases. For variant level QC, extremely rare variants with minor allele frequency *<* 0.05 were removed (PLINK−− maf). For Regenie step 1 model fitting only, we constructed an LD pruned subset of variants (PLINK −− indep-pairwise 50 5 0.2); step 2 used the full QC-passed imputed set. Finally, the genome build was lifted over from hg19 to hg38 with rtracklayer package^9^ in R.

### Weighted Gene Co-expression Network Analysis

We performed cross sectional clustering using Weighted Gene Co-expression Network Analysis (WGCNA)^5^ package in R. The data were partitioned by visit, and the standard WGCNA pipeline was applied separately to each visit-specific dataset to identify modules of co-expressed proteins per visit. Sample outliers were assessed for each visit independently using two methods: hierarchical clustering and principal component analysis. Any sample identified as an outlier at any visit was removed from the datasets of the other visits to maintain a consistent sample set across visits. After this step, each visit had 757 samples and 7288 human proteins.

All WGCNA analyses were performed using a soft-threshold power of 18, biweight midcorrelation (corType = “bicor”), a signed network and TOM (networkType = “signed”, TOMType = “signed”), maxPOutliers = 0.07, reassignThreshold = 0, pamRespectsDendro = TRUE, and a module merge height of 0.25 (mergeCutHeight = 0.25) (Supplementary Figure 3). By default, WGCNA labels modules based on the size: i.e., the largest module - turquoise, second largest - blue, third largest - brown, etc. To have consistent module labels across visits, we relabeled the modules of each visit using the WGCNA function *matchLabels*, which examines the overlap of proteins between modules at different visits to re-label them so that modules with the greatest overlap share the same label.

Protein module membership (kME) was calculated for all proteins across all modules and visits using signedKME function in WGCNA package. It measures a protein’s correlation with its corresponding module eigengene, which helps identify the most important proteins, or “hub proteins/genes,” within a protein co-expression module. A high absolute kME value (e.g., > 0.9) indicates a protein is a highly connected hub within that module.

To evaluate the associations of the clinical phenotypes with the protein modules for each visit, we calculated Pearson correlations between the modules eigengenes (first principal component of each module) and clinical variables, and visualised the associations as a heatmap using the CorLevelPlot function from CorLevelPlot^10^ package in R.

#### Consensus Weighted Co-expression Network

A consensus weighted co-expression network was constructed using the WGCNA^5^ package function blockwiseConsensusModules in R to identify consensus modules across the 3 visits. The parameters used were the same as those used in cross-sectional WGCNA. If the consensus modules overlap with the cross sectional WGCNA modules for each visit, then this would serve as a method validation of the robustness and stability of the cross sectional WGCNA modules for each visit.

### Module Preservation Analysis

#### Cross-sectional wgcna module preservation across visits

To assess the stability of the protein modules across visits, we used the modulePreservation function from WGCNA^5^ R package. This function compares modules from a *reference* set and a *test* set by calculating multiple preservation statistics, followed by an aggregated score, *Z*_*summary*_. In doing so, it uses only the common genes across the two sets. One important parameter in the function is the number of permutations used to calculate the validity of the preservation statistic. We used 1000 permutations and obtained *Z*_*summary*_ score per module for each pair of reference and test, where once, the reference set pertained to V1 and the test set to V2 and V3 and, then V2 was the reference set and V3 was the test set. A *Z*_*summary*_ *<* 2 indicates that there is no evidence of module preservation, 2 *< Z*_*summary*_ *<* 10 indicates weak to moderate evidence of module preservation, and *Z*_*summary*_ *>* 10 indicates strong evidence of module preservation^6^. Furthermore, to assess the temporal stability of the modules, we created a Sankey flow plot to visualize the flow of proteins across modules across visits.

#### Validation

Replication of the module preservation analysis findings from the TP cohort was sought in the Global Neurodegeneration Proteomic Consortium (GNPC)^11^ (v1.3) with samples from multiple independent contributing sites that contributed >10 samples. We selected baseline *Citrate* and *EDTA* plasma samples from individuals with a clinical diagnosis of PD, excluding samples with undefined age or sex. This resulted in 440 samples from three independent contributors, including the PPMI cohort, which were subsequently corrected for contributor-related batch effects (using *combat*^12^) prior to analysis.

### Pathway and Cell Enrichment Analysis

To evaluate the biological relevance of the modules at each visit, we performed pathway enrichment analysis using KEGG database. Each module’s proteins were tested for enrichment against a background of all 7,288 proteins measured using the compareCluster function of clusterProfiler^13^ package in R. The significantly enriched pathways for each module were visualized using a faceted dot plot.

To evaluate the cellular context of the modules at each visit, we performed brain and brain-associated cell-type enrichment analysis using the EWCE^14^ package in R. For each module, human gene symbols mapped from SomaLogic SeqIds were tested against mouse cortex and hypothalamus single-cell RNA-seq data (as reference) using a bootstrap permutation procedure (10,000 permutations; annotation level 1), with cross-species settings appropriate for human gene lists and a mouse reference. Significant cell-type enrichments (FDR-adjusted) were collated across visits and visualized as faceted dot plots by module and cell type.

To identify tissue and cell-type specificity of protein co-expression modules across the three study visits, we performed enrichment analysis using the enrichR^15^ package in R. For each non-grey module at each visit, we queried gene symbols against four tissue/cell-type databases: ARCHS4_Tissues^16^, Allen_Brain_Atlas_10x_scRNA_2021, Allen_Brain_Atlas_Up^17^, and Allen_Brain_Atlas_Down^17^. Enrichment significance was determined using Fisher’s exact test with Benjamini–Hochberg correction for multiple testing. Terms with adjusted *P*-value *<* 0.05 were considered significantly enriched and visualized as faceted dot plots by module and tissue/cell type.

### pQTL Analysis

We hypothesised that proteins whose abundance levels influenced by *cis* acting variants are likely to be enriched in WGCNA modules and potentially act as *hub* proteins within them. We used a baseline protein 4K profile measured on Somalogic platform (4006 proteins) for this analysis.

To establish protein-SNP associations, Regenie^18^ was used. In the first step, a whole-genome regression model was fitted to the LD-pruned subset of genotypes using leave-one-chromosome-out (LOCO) predictions. In the second step, this model was used to rapidly test for associations in every protein-SNP pair from the MAF-filtered full genotype set (not LD pruned). The covariates used in the genome-wide regression model include age, sex, and LEDD (Levodopa Equivalent Daily Dose), together with the first 20 genetic PCs (ancestry) and the first 20 proteomic PCs (technical/batch). Multiple testing correction was done using FDR across all *cis*-signals, and a protein was declared significant if it had at least one variant achieving *q <* 0.05.

### Software and Reproducibility

All the analyses that we performed were executed using the R version 4.3.2 (2023-10-31). Key packages used were WGCNA^5^ (v 1.72-5) for cross sectional clustering, sva^8^ (v 3.50.0) for batch correction, clusterProfiler^13^ (v 4.10.1) for pathway enrichment, EWCE^14^ (v 1.10.2) for cell-type enrichment, enrichR^15^ (v 3.4) for tissue/cell type enrichment, and PLINK^19^ (v1.9) and regenie^18^ (v3.2) for the pQTL analysis.

## Discussion

Using a deeply phenotyped dataset we have generated three robust mechanistic modules of proteins prioritized by clinical relevance. These results demonstrate the application of clustering methods to generate informative modules of disease mechanism. We checked if these clusters were stable and preserved across the visits, using module preservation analysis. We also performed a method validation by performing consensus clustering on the three visit data. We subsequently identified pathways, tissue types and cell types enriched in all modules across visits. We further performed pQTL analysis using genes from the core module pathways. Finally we validated the modules on an independent dataset demonstrating module preservation in combined dataset from three cohorts.

We observed five protein modules with functional enrichment (Figure 3). The blue (signalling) module was associated with quality of life scores for Visit 1 suggesting baseline line proteomic changes have a stronger phenotype relationship for this module (Figure 2). The brown (metabolic) module was associated to semantic fluency at Visit 1 but disease duration and age at Visit 3. This may reflect the underlying heterogeneity of the clinical phenotypes despite the consistency of the protein modules. The red (cellular processes) module, despite changes in protein expression, was consistently associated with the age and disease duration and also showed a correlation with MoCA, semantic fluency and RBD at Visit 3 pointing to a cognitive component. Of note in the analysis of clinical phenotypes is the dominance of aging as a key phenotype (Figure 2) . Given the dominance of age as the primary risk factor for PD this was an expected result, however as it is not a distinct pathology and has strong associations to different models of expression, it was not possible to normalise the data without reducing relevant disease and progression variation.

As with a previous study using the TP and OPDC observational cohorts, the largest turquoise module was enriched for inflammatory proteins and pathways^20^. This suggests the protein expression module is robust to Somalogic assay version. However, although the module was preserved across time points it was not found in the external validation sets indicating a potential cohort specific data module or protein set impacted by differences between the plasma and serum matrices, likely given the sensitive nature of inflammatory assays.

Inflammatory pathways in particular were common in two modules and co-expression within both these modules was consistent between time points. The blue (signaling) module was enriched for T cell receptor and chemokine signaling pathways. There is clear evidence of the association between inflammation and PD in other studies, however, what is not always so obvious is the number of associated changes relating to this mechanism. These pathways are not unique to PD and therefore biomarkers relevant to other neurodegenerative diseases could act to inform on the inflammatory subtypes.

Validation of the results was key in demonstrating the robustness of the other modules. Using the GNPC, PD cases showed strong preservation for the brown (metabolic) and blue (signaling) modules (Figure 5B). This indicates that the co-expression of the component proteins is common in other cohorts and a feature of disease. A striking feature of this analysis approach was the robustness of the clusters over time. Cluster preservation between the three key time points was strong with even smaller modules showing weak evidence of preservation (zsummary<2).

Replication of clusters remains a core problem of predicting disease subtypes. Although we report preserved protein modules, other studies report alternative subtypes. Using data from the PPMI study and deep learning algorithms, Zhang et al^21^ defined three subtypes but also showed the complexity of combining clinical and biomarker data, in particular, that baseline severities are not predictive of progression rates. Data-driven stratification of the TP cohort by clinical features has led to the identification of four subtypes^22^. This demonstrates the heterogeneity of PD in the cohort patients in clinical features in addition to the protein heterogeneity highlighted in our own analysis. Using Bayesian mixed model approaches to combine genetics and clinical progression data led to the identification of three AD-like disease axes^23^. Here, we use a single modality and define our module subtypes by protein pathophysiology before utilizing other measures (clinical and genetic) to confirm and prioritize output. Validation has demonstrated that these protein modules can be used in other cohorts.

The overlap of pQTL proteins with module proteins suggests that genetic predisposition may affect the key machinery of the pathways in *brown, blue*, and *black* modules. Specifically, the metabolic pathways, signaling pathways, and infection and disease related pathways enriched in *brown, blue*, and *black* modules, respectively, that share proteins with the pQTL analysis hits are likely to play a role in the progression of the disease in PD patients.

There are a number of important considerations and potential limitations to the study. Firstly, protein measures in the TP cohort were generated from serum not CSF, meaning the modules are representative of combined peripheral and brain expression changes. Although our data has shown independent modules for peripheral protein’s expression it still makes interpretation of a full body expression signature complex. Results will strengthen outcomes for prognosis biomarker detection in disease subtypes but screening for potential drug target should be implemented on a module and pathway level. Inconsistency between high throughput platforms makes it difficult to replicate between studies. Here, we compare data from Somalogic measured assays which, although gives us access to more proteins, limits validation on OLINK due to the lack of overlapping assays in some cases. Our replication in Somalogic generated datasets ensures module reproducibility by using comparable aptamer assays, protein isoforms and blood sample matrix. Secondly, for the pQTL analysis, we had a moderate sample size ( ∼ 800) and we focused only on *cis* signals as a result. Furthermore, despite extensive QC and covariate adjustments, site-specific artefacts remain possible. Therefore, the findings should be validated in a larger independent cohort, which can also help detect *trans* signals without being limited by statistical power. Thirdly, the TP cohort is a UK based cohort and although it is geographically and socio-economically representative of the UK, it lacks a broader ethnic diversity. However, the cohorts incorporated in validation as part of the GNPC are from cohorts outside the UK and includes US populations (such as PPMI) and broader European representation. This demonstrates the applicability across different cohorts of these modules.

In summary, we have shared a set of novel robust modules of proteomic subtype for PD which are preserved longitudinally and across cohorts. Detected in blood samples although expressed in brain, the validated protein modules (signaling and metabolic) will be utilized to better understand disease mechanism and progression and establish potential markers to group patients by pathology using proteomic blood assays.

## Acknowledgments

The authors would like to thank the patients who contributed to the studies used in the manuscript. L.W is funded by Alzheimer’s Research UK (ARUK-RF2020A-005, ARUK-SRF2023B-007). R.M and B.E are funded by Michael J. Fox Foundation (MJFF-022845). H.M. receives Grants from Parkinson’s UK, Cure Parkinson’s Trust, PSP Association, Medical Research Council, Michael J Fox Foundation and NIHR.

## Author contributions statement

L.W and D.G. conceived the study, R.M and B.E were responsible for methodology and formal analysis, R.M., R.R. and S.G. data curation and cleaning, R.M, B.E and L.W were responsible for original draft preparation. All authors reviewed the manuscript.

## Additional information

Proteomic data described in this study is available on application to GNPC.

