## Supplementary Material for "Longitudinal Proteomic Profiling Defines Robust Molecular Subtypes Underlying the Heterogeneity of Parkinson’s Disease"

**Figure S1. Consensus WGCNA protein module pathway enrichments**

| 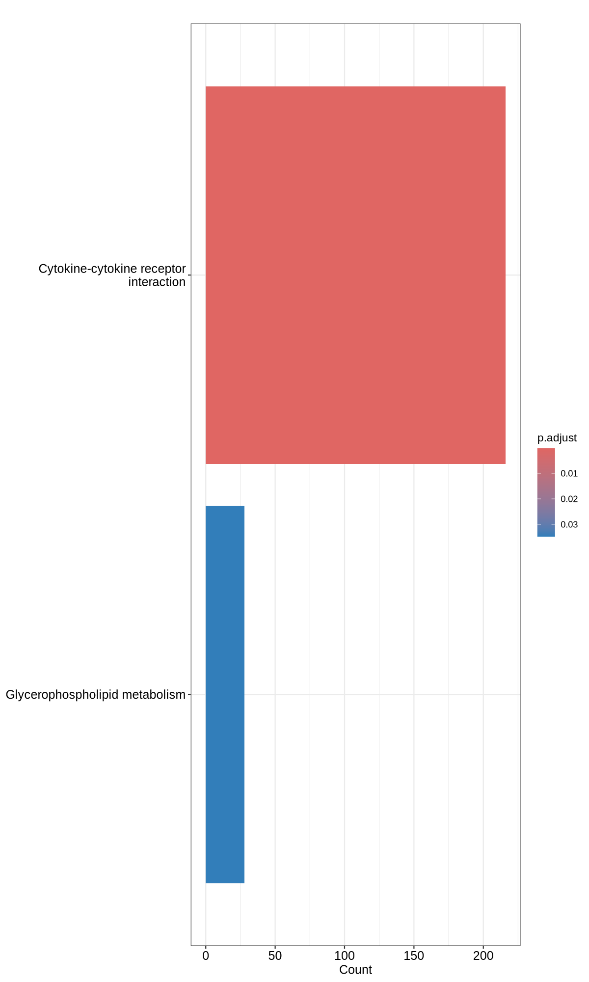a | 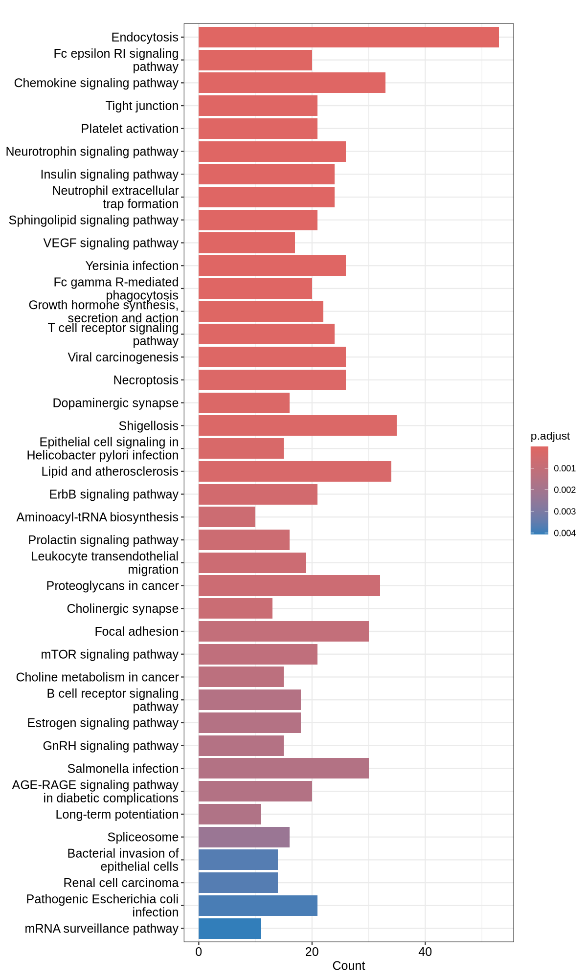b |
| --- | --- |
| 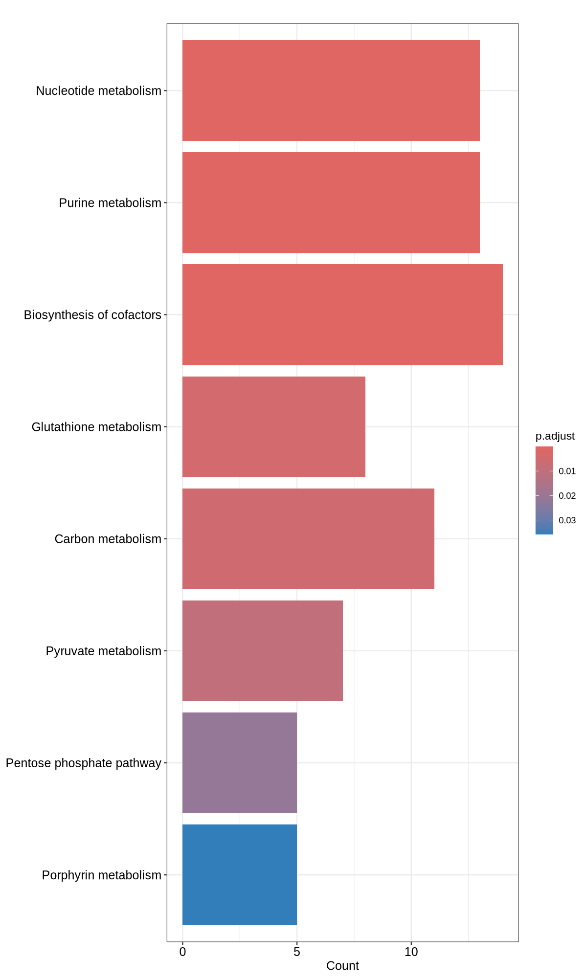c | 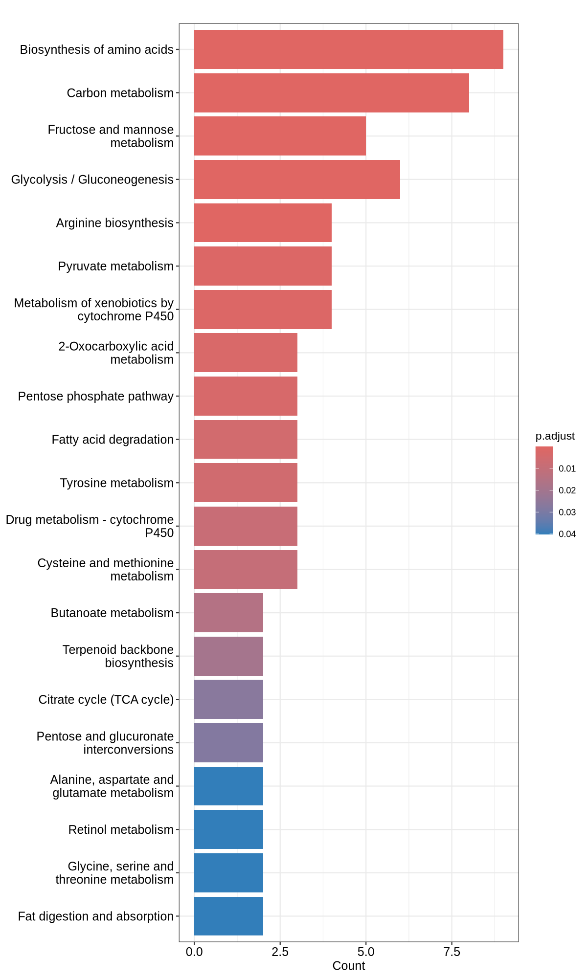d |
| 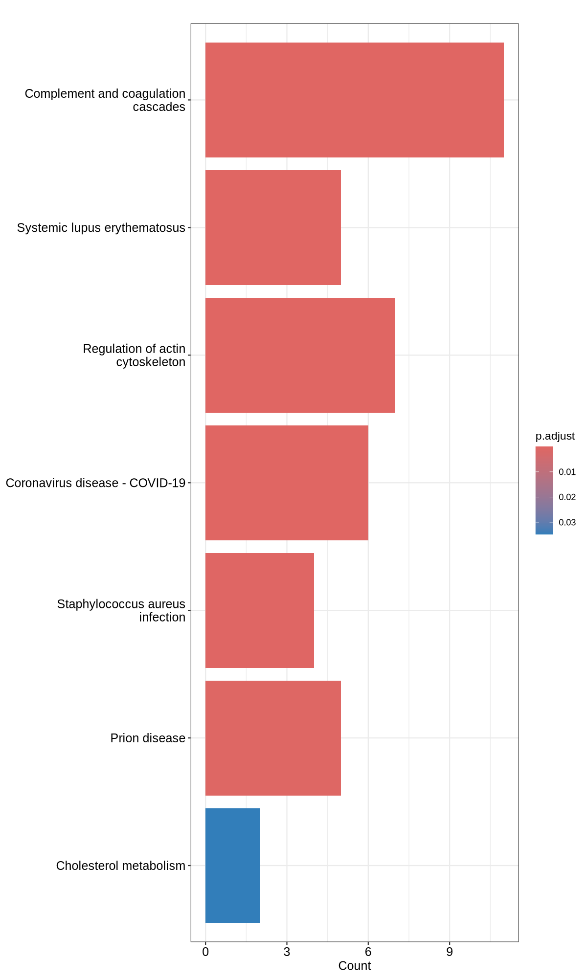e | |

(a) Turquoise module (b) Blue module (c) Brown module (d) Red module (e) Black module. Top 20-40 significantly enriched pathways are shown here

**Figure S2 (a). Protein module preservation across the three visits using medianRank for the discovery cohort.**

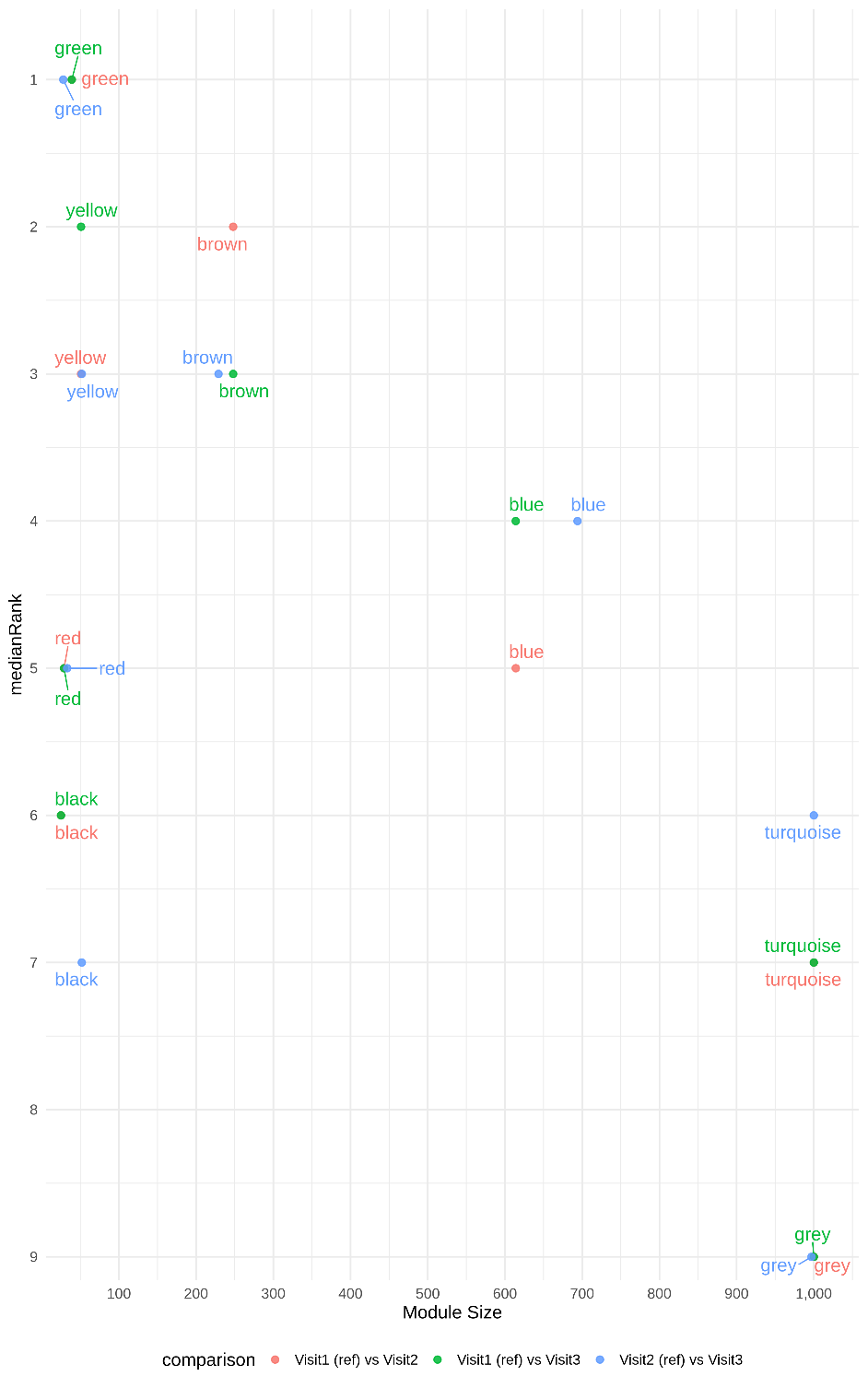

medianRank is a composite statistic for module preservation which considers all modules on an equal footing irrespective of module size. Low medianRank (eg 1 or 2) for a module means the module is strongly preserved

**Figure S2 (b). Protein module preservation across the three visits using medianRank for the validation cohorts (GNPC).**

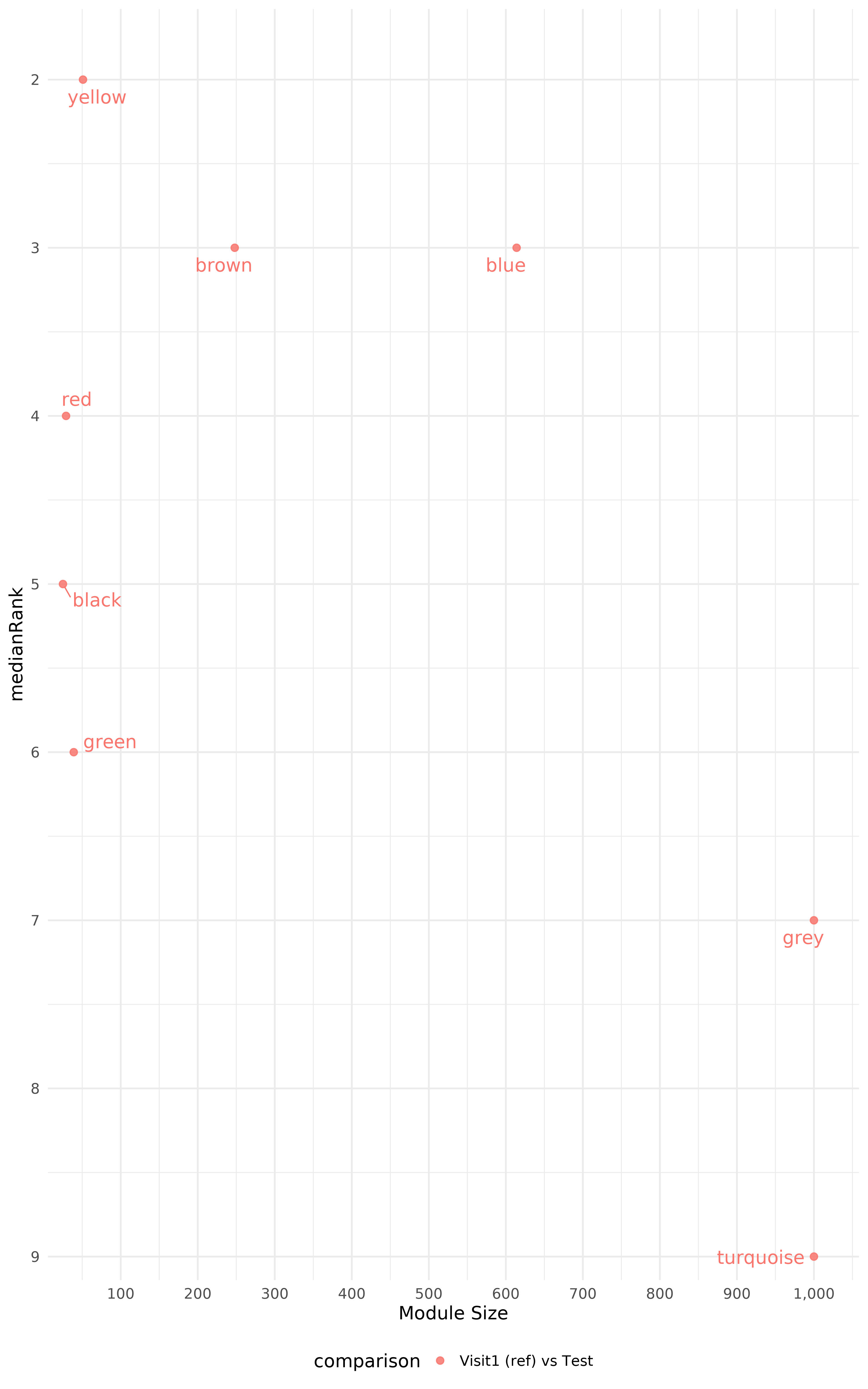

**Figure S3. Soft-thresholding power selection and module detection for weighted gene co-expression network analysis (WGCNA).**

| 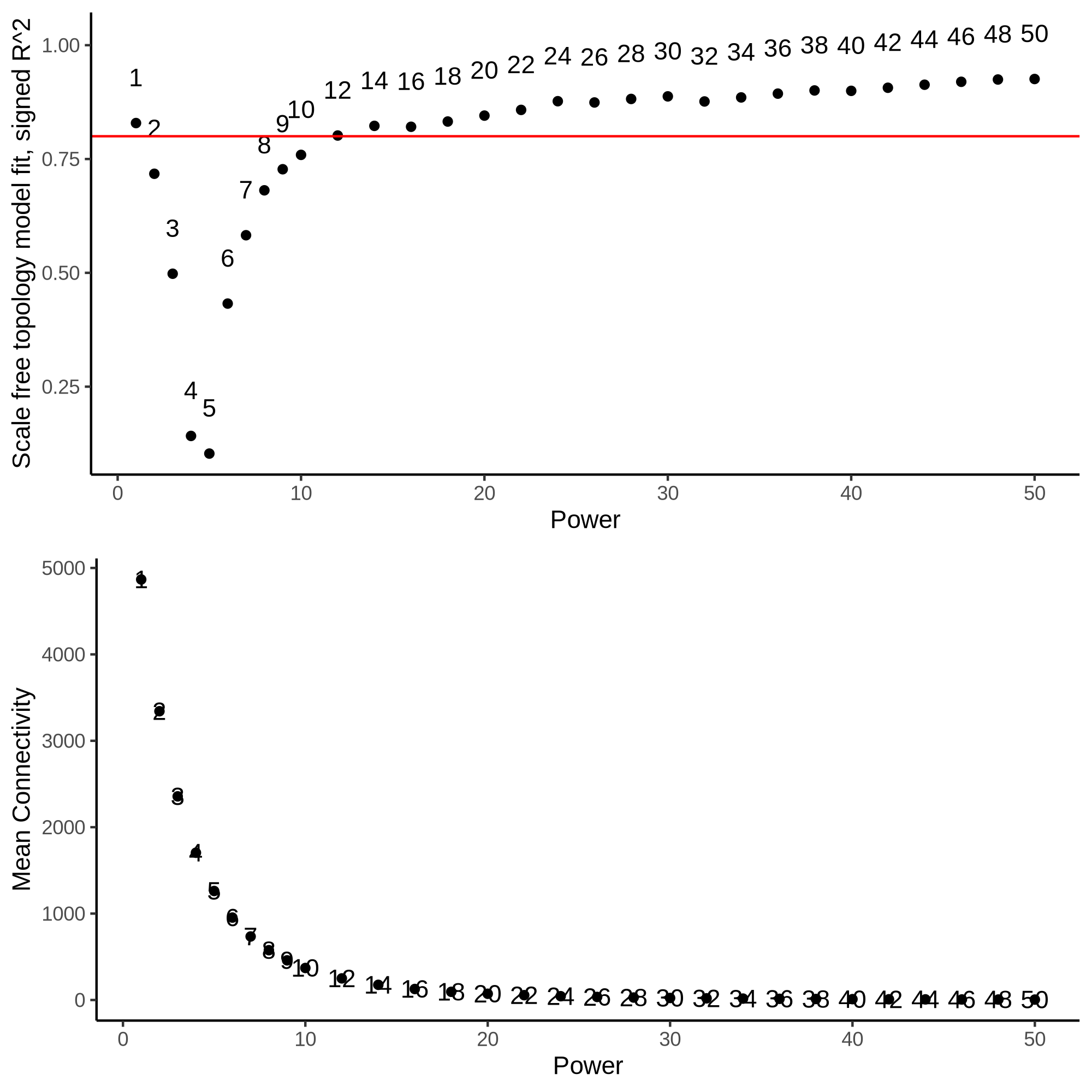a | 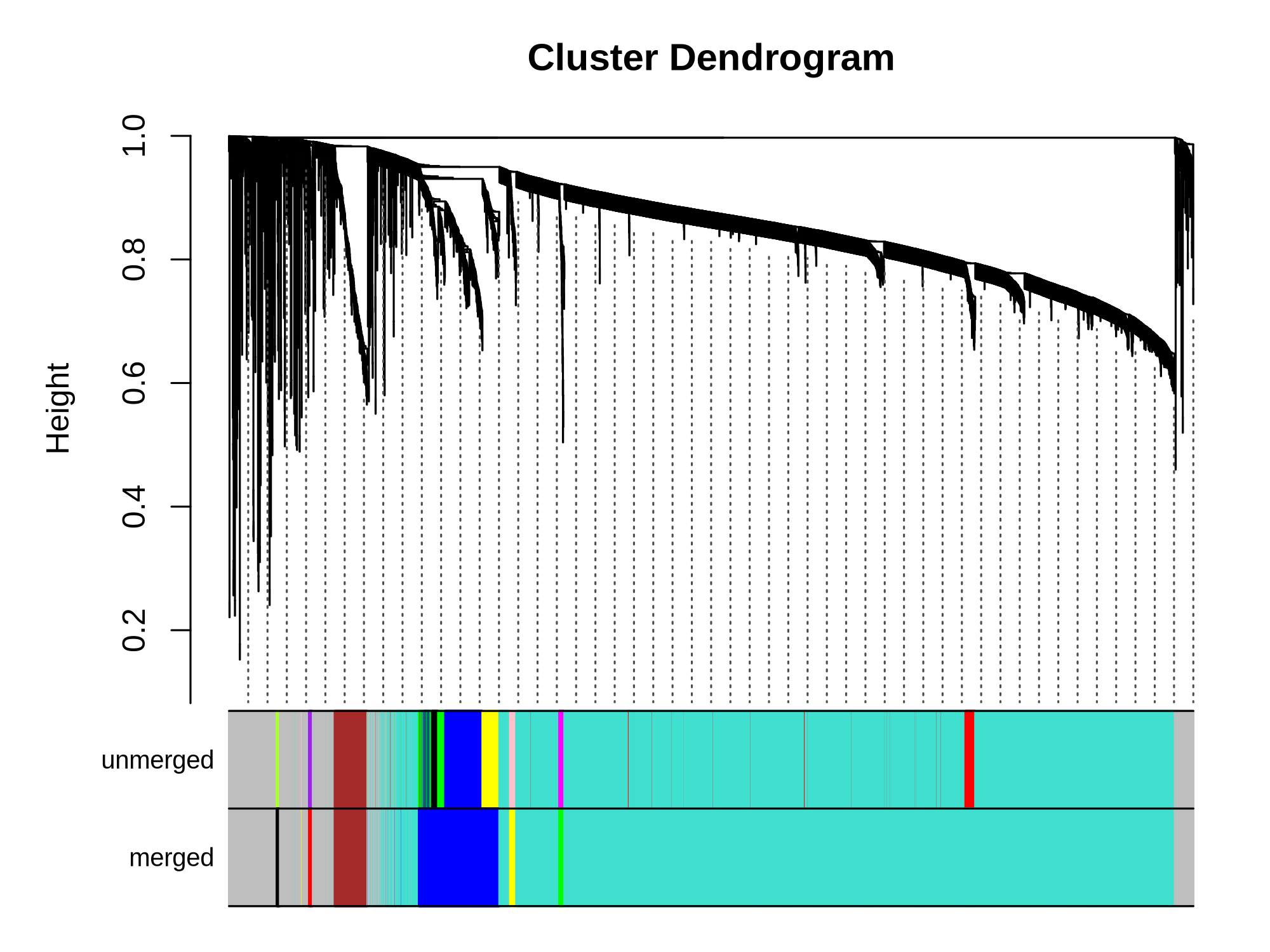b |
| --- | --- |
| 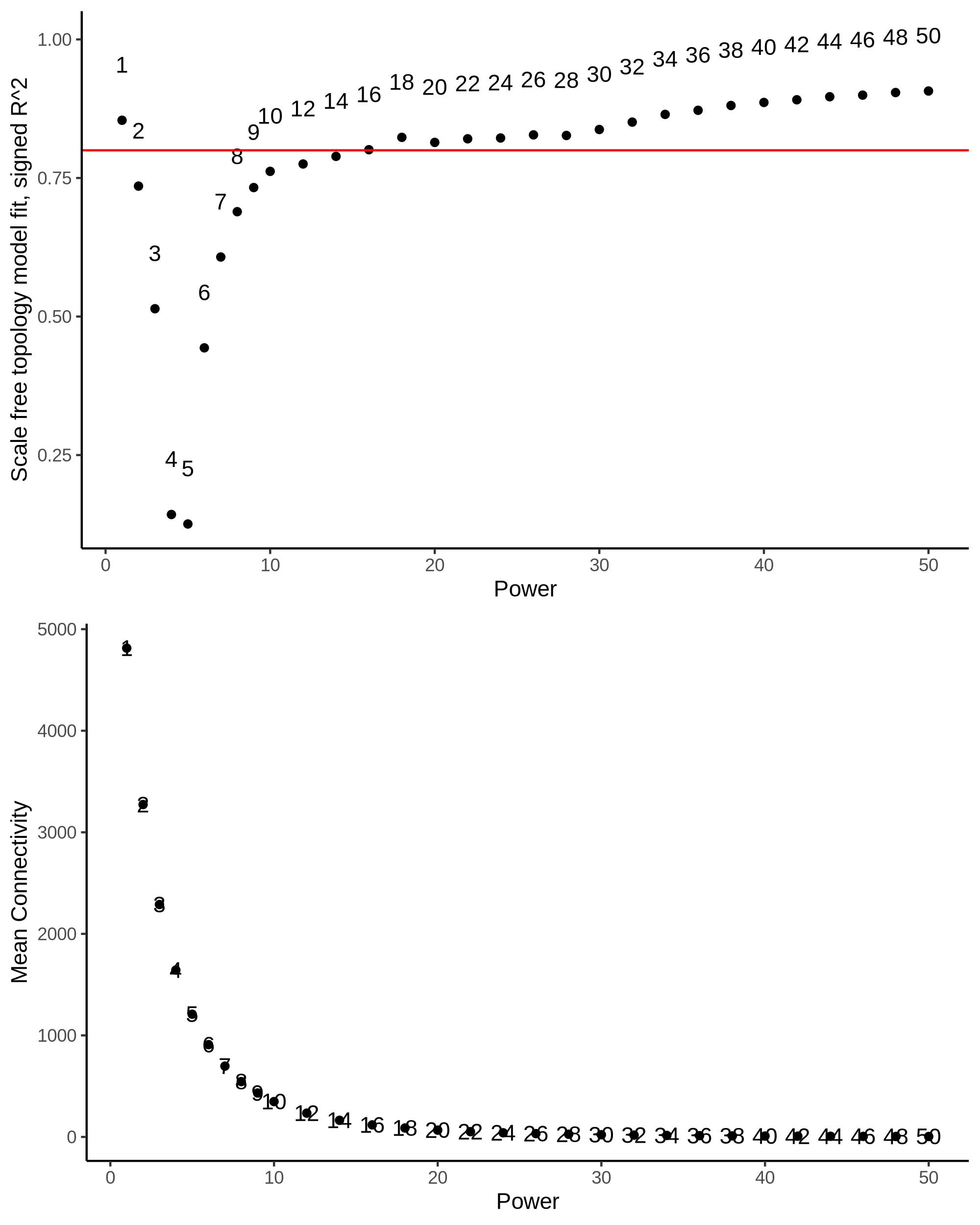  c | 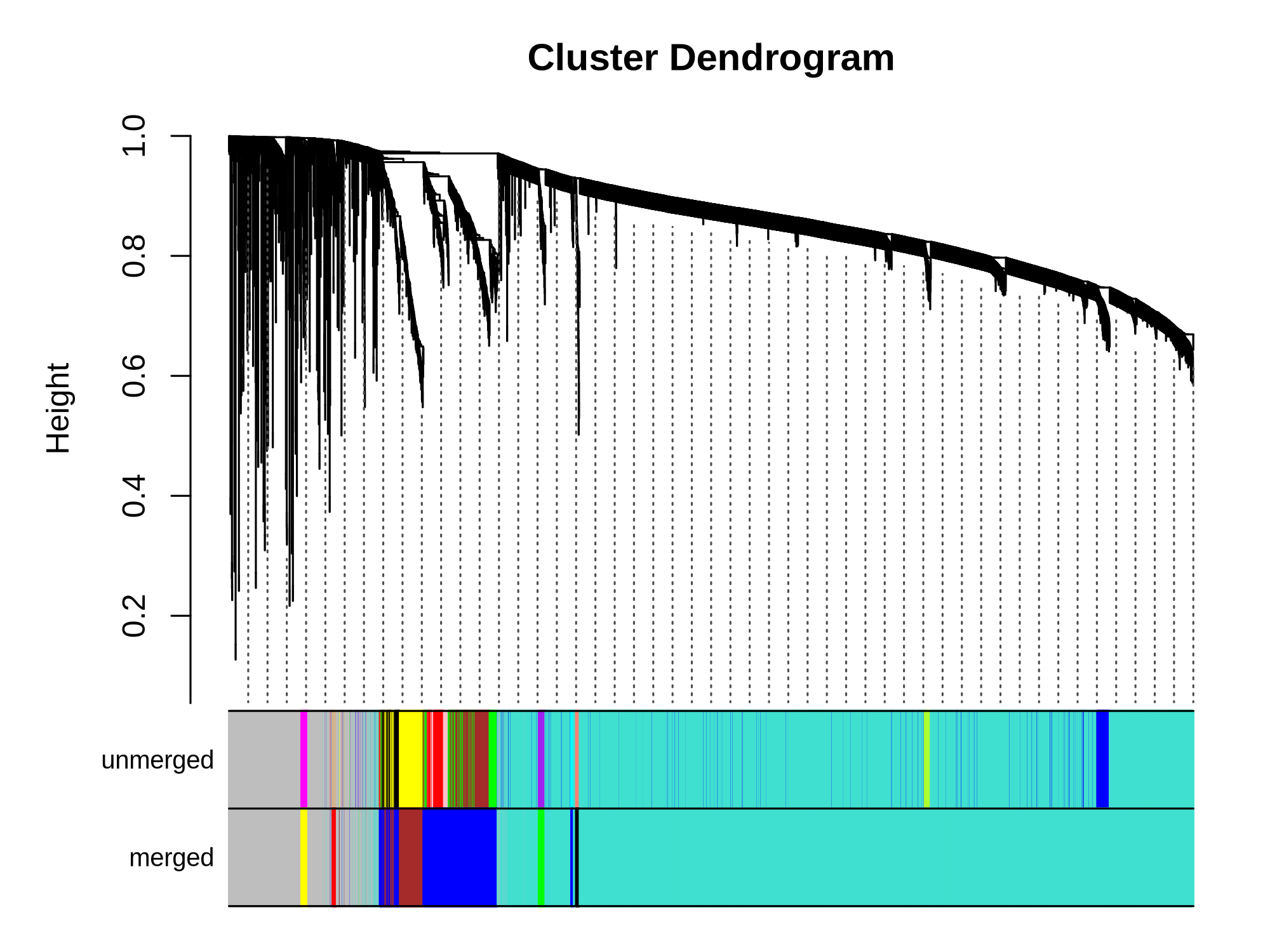d |
| 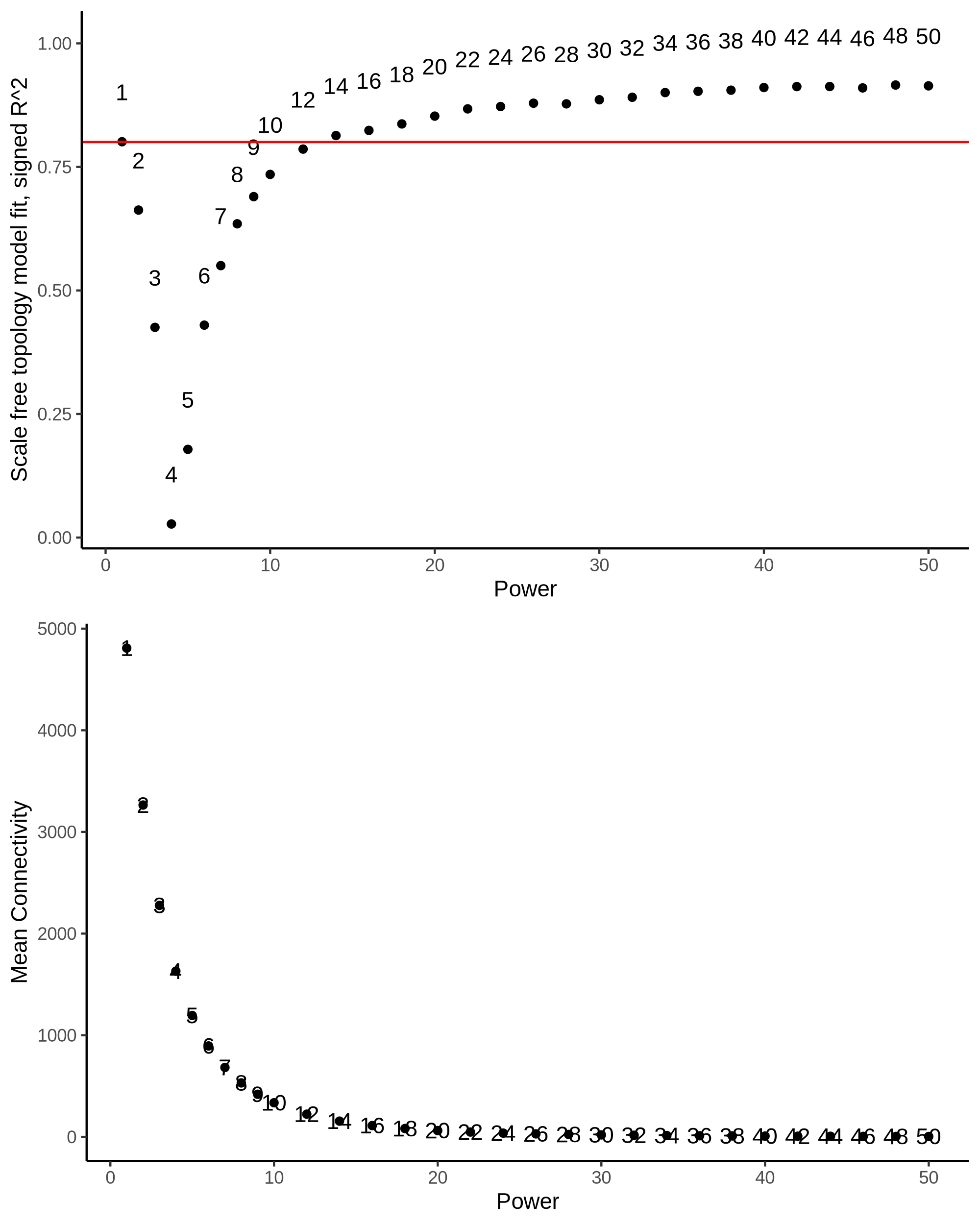  e | 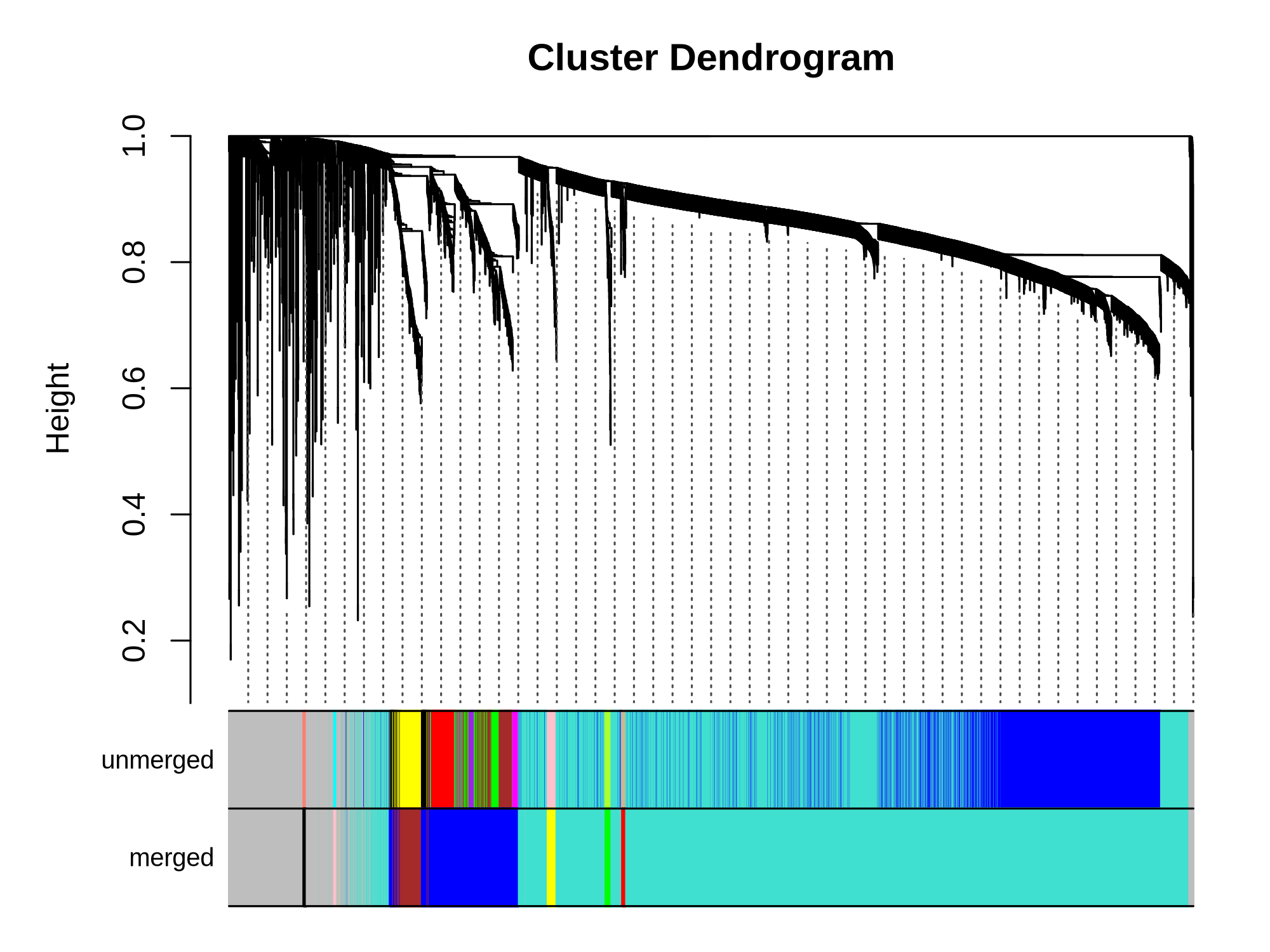f |

(A, C, E) Soft-thresholding power selection for Visit 1, Visit 2, and Visit 3, respectively. Top panels show scale-free topology model fit (signed R²) across different soft-thresholding powers. The red horizontal line indicates the recommended threshold of R² = 0.8. Bottom panels show mean connectivity across the network for each power. The selected power balances high scale-free topology fit (R² > 0.8) with reasonable mean connectivity. (B, D, F) Hierarchical clustering dendrograms with module assignments for Visit 1, Visit 2, and Visit 3, respectively. The dendrogram shows hierarchical clustering of proteins based on topological overlap. Two color bars below each dendrogram represent module assignments before merging (unmerged) and after merging similar modules. Each color represents a distinct co-expression module.
